# Integrative genome-scale metabolic modeling reveals versatile metabolic strategies for methane utilization in *Methylomicrobium album* BG8

**DOI:** 10.1101/2021.03.21.436352

**Authors:** Juan C. Villada, Maria F. Duran, Chee Kent Lim, Lisa Y. Stein, Patrick K. H. Lee

**Affiliations:** School of Energy and Environment, City University of Hong Kong, Hong Kong SAR; Department of Biological Sciences, University of Alberta, Edmonton, AB, Canada

**Keywords:** Methanotroph, methane oxidation, systems biology, integrative modeling, flux balance analysis, fermentation

## Abstract

*Methylomicrobium album* BG8 is an aerobic methanotrophic bacterium that can mitigate environmental methane emission, and is a promising microbial cell factory for the conversion of methane to value-added chemicals. However, the lack of a genome-scale metabolic model (GEM) of *M. album* BG8 has hindered the development of systems biology and metabolic engineering of this methanotroph. To fill this gap, a high-quality GEM was constructed to facilitate a system-level understanding on the biochemistry of *M. album* BG8. Next, experimental time-series growth and exometabolomics data were integrated into the model to generate context-specific GEMs. Flux balance analysis (FBA) constrained with experimental data derived from varying levels of methane, oxygen, and biomass were used to model the metabolism of *M. album* BG8 and investigate the metabolic states that promote the production of biomass and the excretion of carbon dioxide, formate, and acetate. The experimental and modeling results indicated that the system-level metabolic functions of *M. album* BG8 require a ratio > 1:1 between the oxygen and methane specific uptake rates for optimal growth. Integrative modeling revealed that at a high ratio of oxygen-to-methane uptake flux, carbon dioxide and formate were the preferred excreted compounds; at lower ratios, however, acetate accounted for a larger fraction of the total excreted flux. The results of this study reveal a trade-off between biomass production and organic compound excretion and provide evidence that this trade-off is linked to the ratio between the oxygen and methane specific uptake rates.

---

Anthropogenic activities have led to significant increases in atmospheric methane^1^, which contributes to climate change and perturbs the global carbon cycle ^2^. However, methane derived from renewable sources is an attractive substrate in the production of value-added products ^3–5^, and methane-conversion processes represent a promising trend in bioindustry ^6,7^. Methanotrophic bacteria utilize methane as their source of carbon and energy, and these microorganisms have become increasingly important in the biomanufacturing of valuable chemical compounds ^4,8–10^. Although methanotrophic species are metabolically active under both aerobic and anaerobic conditions ^11–14^, considerations of cost, sustainability, and environmental impact have led to a preference for aerobic methane-oxidizing bacteria in large-scale biorefining applications ^15,16^.

*Methylomicrobium album* BG8 (formerly known as *Methylobacter albus*, *Methylomonas albus*, or *Methylomonas alba*) is an obligate aerobic, gram-negative, gammaproteobacterial methanotroph that uses methane or methanol as its sole source of carbon and energy ^17^. A DNA-DNA hybridization study revealed high levels of similarity between its genome and those of *Methylomicrobium agile* ATCC 35068 (99.16%), *Methylotuvimicrobium alcaliphilum* 20Z (75.69%), and *Methylotuvimicrobium buryatense* 5G (76.64%) ^18^. Recent phylogenomic analyses based on the average amino acid identity and average nucleotide identity have shown that *M. album* BG8 is also closely related to *Methylomicrobium* (formerly *Methylosarcina*) *lacus* LW14 ^19^. *M. album* BG8 has been isolated from swampy soils and freshwater sediments ^18,20^ and has been widely studied due to its importance in environmental microbiology for bioremediation of different environmental pollutants ^21–23^. Through recent physiological and omics analyses ^24,25^, *M. album* BG8 has also been identified as a promising microbial cell factory for applications in the methane-based biotechnology industry ^25^.

The metabolic engineering of methanotrophs has enabled advances in the biotechnology of methane conversion to produce succinate ^26^, 3-hydroxypropionic acid ^27^, 2,3-butanediol ^28^, putrescine ^9^, *α*-humulene ^29^, cadaverine ^30^, lysine ^30^, shinorine ^31^, and acetoin ^31^. These laboratory achievements have been aided by the results of genome-scale metabolic models (GEM)-based simulations, which are used to enhance the system-level understanding of methanotrophy. Thus far, GEMs of nine methanotrophic species have been constructed, including three type I species [Gammaproteobacteria; *Methylotuvimicrobium buryatense* 5G(B1) ^32^, *Methylotuvimicrobium alcaliphilum* 20Z ^33^, and *Methylococcus capsulatus* Bath ^34^,^35^] and six type II species [Alphaproteobacteria; *Methylocystis hirsuta* ^36^, *Methylocystis sp*. SC_2_^36^, *Methylocystis sp*. SB_2_^36^, *Methylocystis parvus* OBBP ^37^, *Methylocella silvestris* ^38^, and *Methylosinus trichosporium* OB3b ^39^]. Although some of these GEMs have been validated using growth yields, methane and oxygen specific uptake rates ^36^, transcriptomics data ^32^, or enzyme kinetics ^33^, most do not contain integrated experimental data.

Despite the enormous potential of *M. album* BG8 as a tool in environmental and industrial biotechnology, no GEM of this methanotroph has been developed, and therefore, an integrative system-level understanding of methanotrophy in this strain remains lacking. In this study, a high-quality GEM of *M. album* BG8 was constructed by stringently following well-established systems biology protocols ^40,41^. Furthermore, an integrative modeling framework was applied, wherein experimental time-series growth and exometabolomics data collected under different initial methane and oxygen headspace percentages and biomass concentrations were integrated with the initial GEM to construct context-specific GEMs (csGEMs). Subsequently, the metabolic states that promote biomass production and carbon dioxide, formate, and acetate excretion were identified through a flux balance analysis (FBA). The study findings provide novel insight into the metabolic versatility of *M. album* BG8 and highlight the trade-off between biomass production and organic compound excretion.

## Results

### Relationship between optical density (OD) and dry cell weight (DCW)

The primary objective of this study was to analyze the metabolism of *M. album* BG8 at a cellular system level, and the required metrics and model parameters depend on DCW-based data. Hence, a linear regression model was established between *l*^−1^ and OD_600_ to calculate the DCW from a measured OD_600_ value. The model with a y-intercept at the origin produced the best fit (Fig. S1), yielding the equation *gDCW l*^−1^ = 0.26 *OD*_600_.

### Excess methane did not favor metabolite production

Aerobic methane oxidation, which is catalyzed by the particulate methane monooxygenase in *M. album* BG8, requires an equimolar ratio of oxygen to methane. We first hypothesized that excess methane in a batch culture of *M. album* BG8 favors growth. To test this hypothesis, we characterized the effects of two initial methane [20% (0.41 mmol) or 45% (1.0 mmol)] and oxygen [20% (0.43 mmo)] headspace ratios on metabolite excretion and biomass production. The time-series profiles in Fig. 1A demonstrate that a notable amount of methane remained in the culture with 45% initial methane, and that the final concentrations of biomass and excreted products were not significantly different between the cultures grown under different headspace ratios (*t*-test, 5% threshold for *P*-value) (Fig. 1A). Contrary to our hypothesis, excess methane significantly reduced (*t*-test, *P* < 0.05) not only the biomass yield but also the oxygen uptake and carbon dioxide and acetate excretion yields (Fig. 1B). Although the formate yield was also reduced, this difference was not statistically significant.

**Figure 1.**
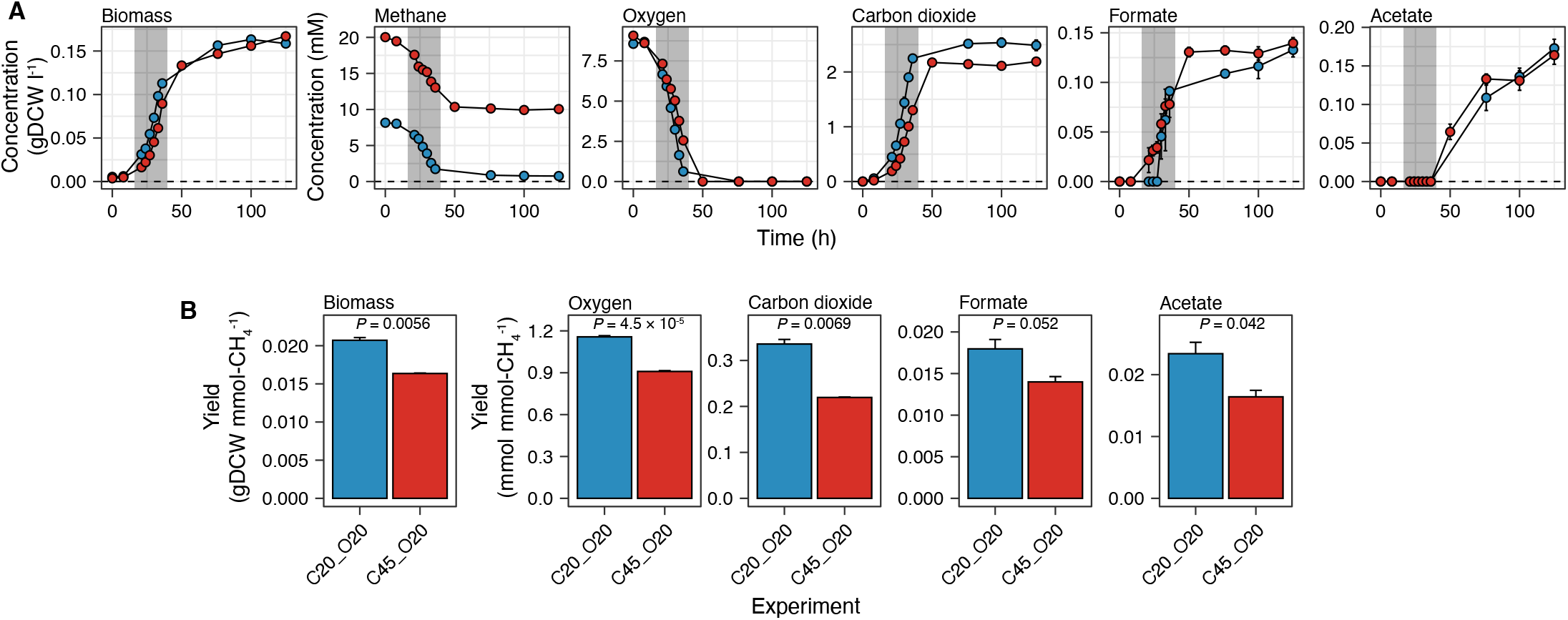
Effects of different initial methane headspace percentages on *Methylomicrobium album* BG8 in culture. **(A)** Time-series concentration profiles of biomass production, compound consumption, and compound excretion. The gray-shaded region indicates the estimated exponential phase (~16 h to 40 h). **(B)** Biomass yields, oxygen uptake, and metabolite excretion throughout the culture period. *P*-values were determined using the two-sided *t*-test. In all panels, the data are shown as the mean values of biological triplicates, and the error bars represent the standard errors. The initial conditions with 20% methane and 20% oxygen (C20_O20) or 45% methane and 20% oxygen (C45_O20) are indicated in blue and red, respectively. In all panels, the same labeling notation is used to denote the experimental conditions, with C representing methane, O representing oxygen, and the following number indicating the initial percentage (v/v) in the headspace.

### Oxygen availability strongly influenced biomass production and organic compound excretion

After determining that excess methane did not favor metabolite excretion, we tested the effects of oxygen availability (range: 5% to 25%) on the metabolism and growth of *M. album* BG8 under 20% methane. The time-series profile in Fig. 2A shows that the initial oxygen headspace percentage significantly affected the dynamics of biomass production and organic compound excretion. The highest concentration of oxygen led to an increased consumption of methane and the highest concentration of excreted organic compounds in culture (Fig. 2A). Interestingly, acetate was only detected after approximately 50 hours in cultures subjected to all conditions except 5% oxygen (~36 h; Fig. 2A). The final carbon dioxide, formate, and acetate concentrations were directly proportional to the initial oxygen headspace percentage (Kruskal-Wallis test, *P* < 0.05; Fig. 2B). Although the final biomass concentration was positively related to the initial oxygen headspace percentage (Kruskal-Wallis test, *P =* 0.011), this variable did not differ substantially between the cultures with 20% and 25% oxygen (Fig. 2B). The carbon dioxide and formate yields and the oxygen uptake were significantly enhanced (Kruskal-Wallis test, *P* < 0.05) by the initial oxygen headspace percentage, whereas the acetate and biomass yields were not significantly affected (*P >* 0.05; Fig. S2). Oxygen was practically depleted from all of the cultures at the end of the experiment (120 h; Fig. 2B).

**Figure 2.**
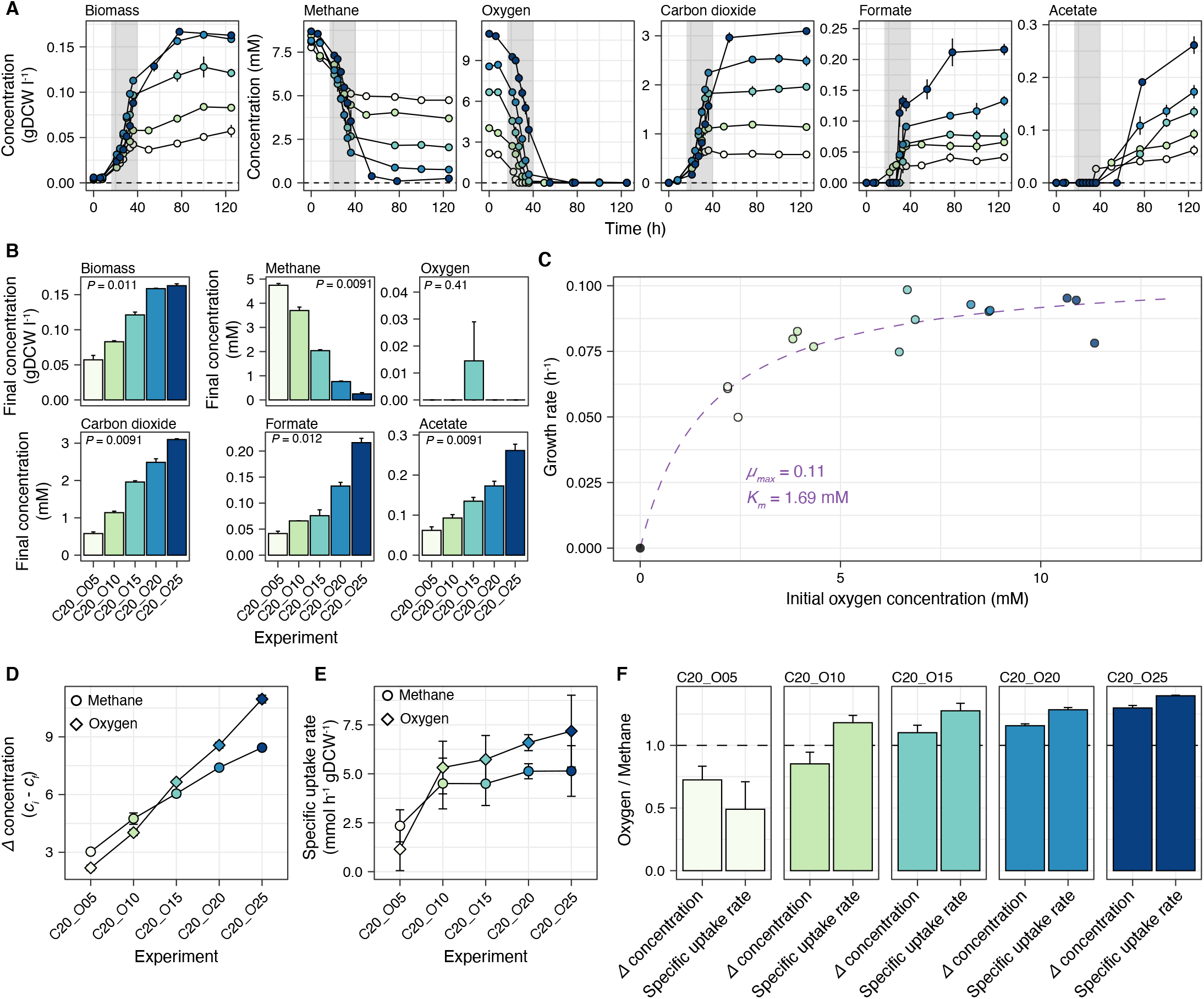
Effects of different initial oxygen headspace percentages on *Methylomicrobium album* BG8 in culture. **(A)** Time-series concentration profiles of biomass production, compound consumption, and compound excretion. The gray-shaded region indicates the estimated exponential phase (~16 h to 40 h). **(B)** The concentrations of biomass, consumed compounds, and excreted compounds at the final time point. (C) Growth kinetics with respect to the initial oxygen concentration. The fitted line represents the Monod kinetic model. **(D)** Changes in methane and oxygen concentrations throughout the culture period. **(E)** Specific methane and oxygen uptake rates in each culture. Error bars represent the upper and lower bounds of the estimate (99% confidence interval). **(F)** Comparison of the ratio between the changes in the concentrations of oxygen and methane with the ratio of oxygen-to-methane specific uptake rates. For panels **A**, **B**, and **D**, the data are shown as the mean values of biological triplicates, and the error bars represent the standard errors. In all panels, the color gradient represents the different initial percentages of oxygen (5% to 25%) in 5% increments with 20% methane.

The growth kinetics of the cultures, assuming oxygen was the only limiting substrate, were determined by fitting a Monod kinetic model, resulting in a maximum growth rate (*μ_max_*) of 0.11 *h*^−1^ [95% confidence interval (CI95%) = 0.09–0.12] and a half-saturation constant (*K_*m*_*) of 1.69 mM (CI95% = 0.78–2.60) (Fig. 2C). The change in oxygen concentration as a function of the initial ratio of oxygen to methane in the culture headspace (i.e., oxygen-to-methane headspace ratio; slope of the oxygen curve) was greater than the change in the methane concentration (i.e., slope of the methane curve) (Fig. 2D), suggesting that the effects of the initial availability of oxygen and methane on the respective proportions of both substrates were mainly mediated by higher oxygen consumption (Fig. 2D). The specific uptake rate of oxygen tended to be higher than that of methane in all experimental conditions except the culture containing the lowest initial oxygen headspace percentage (Fig. 2E). In the cultures with initial oxygen headspace percentages ranging from 10% to 25%, the ratio between the specific uptake rates of oxygen and methane remained similar (1.2:1 to 1.4:1) while the ratios between the changes in the concentrations of oxygen and methane varied from 0.75:1 to 1.3:1 (Fig. 2F). Interestingly, only the culture with the lowest initial oxygen headspace percentage had a low ratio between the specific uptake rates of oxygen and methane (~0.5:1, Fig. 2F) and a noticeably lower rate of specific growth (Fig. 2C).

### Features of the GEM of *M. album* BG8

Given the strong effect of the initial oxygen headspace percentage on the growth rate and metabolic yields, we next aimed to elucidate the metabolic states induced by different levels of oxygen availability in *M. album* BG8. To achieve a system-level understanding of *M. album* BG8 metabolism, we constructed a high-quality GEM that incorporated all of the central metabolic pathways contributing to methane oxidation (Fig. 3) and many other pathways that provide energy and precursors for biomass production in *M. album* BG8 (Fig. 4). The final GEM was established after manually curating the gene-protein-reaction (GPR) associations and biomass compositions in the draft model generated by KBase ^41^, which led to the addition of 16 genes, 33 metabolites, and 181 reactions. The final curated GEM comprises 803 genes, 1,367 metabolites, and 1,358 reactions (Fig. 4A), uses the nutrient parameters described in the Materials and Methods section, and constrains only methane and oxygen uptake fluxes by default (Fig. 4B). Furthermore, the final model purposely includes five blocked reactions (i.e., flux is not allowed) (Fig. 4C), of which four are involved in alternative biomass production and one involves an alternative methanol dehydrogenase, and 171 orphan reactions (i.e., with no GPR associations).

**Figure 3.**
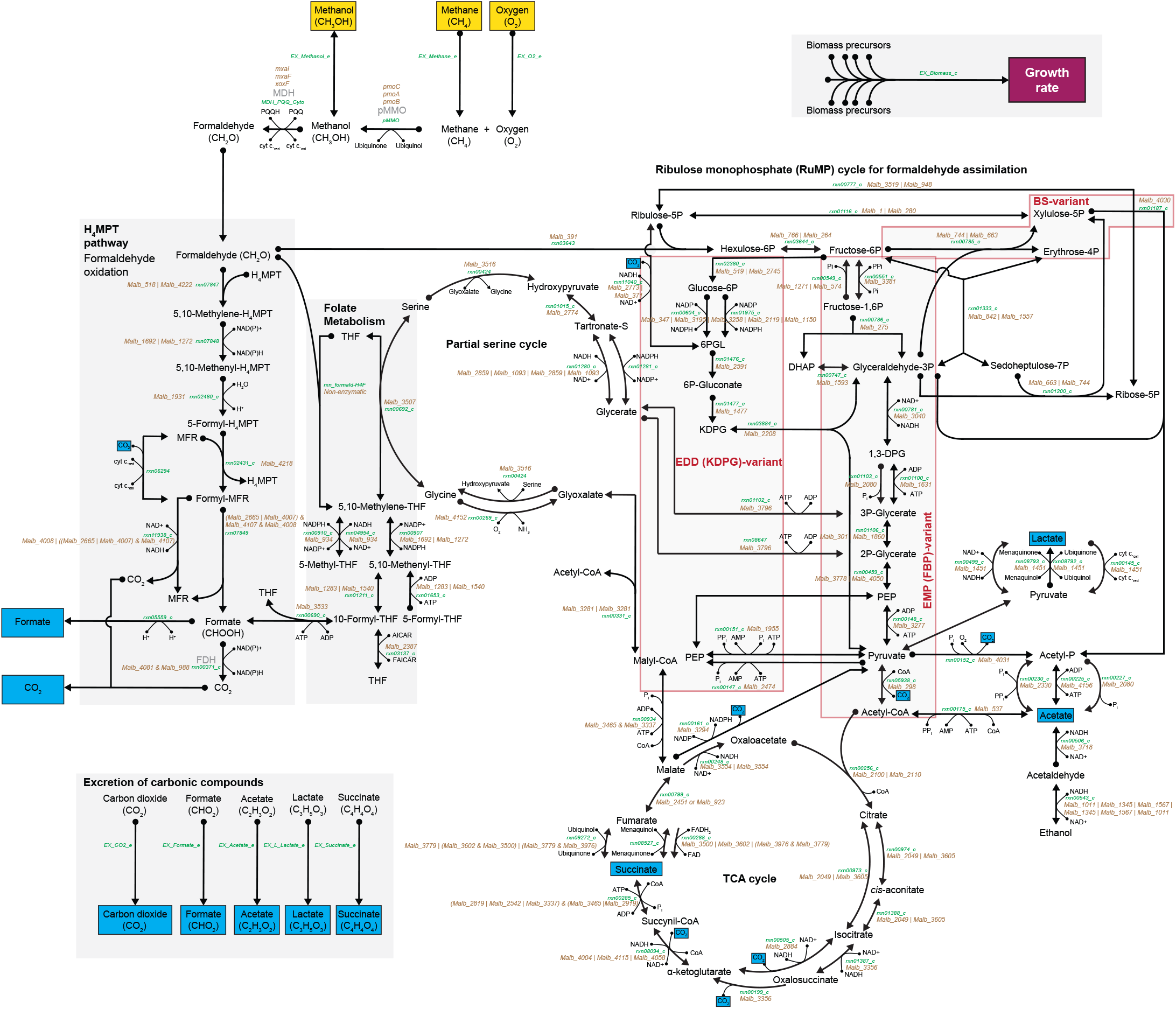
Central methanotrophic metabolic pathways in *Methylomicrobium album* BG8. Metabolites that are taken up are highlighted in yellow, and those excreted are labeled in blue. Metabolite names are shown in black, reaction identification numbers are in green, and the genes associated with each reaction are in brown. The pathway names are bolded. The chemical formula describing the key metabolites involved in the initial methane oxidation and the consumed and excreted metabolites is shown. Non-reversible and reversible reactions, determined according to the thermodynamics constraints of every reaction, are indicated by single-headed and double-headed arrows, respectively.

**Figure 4.**
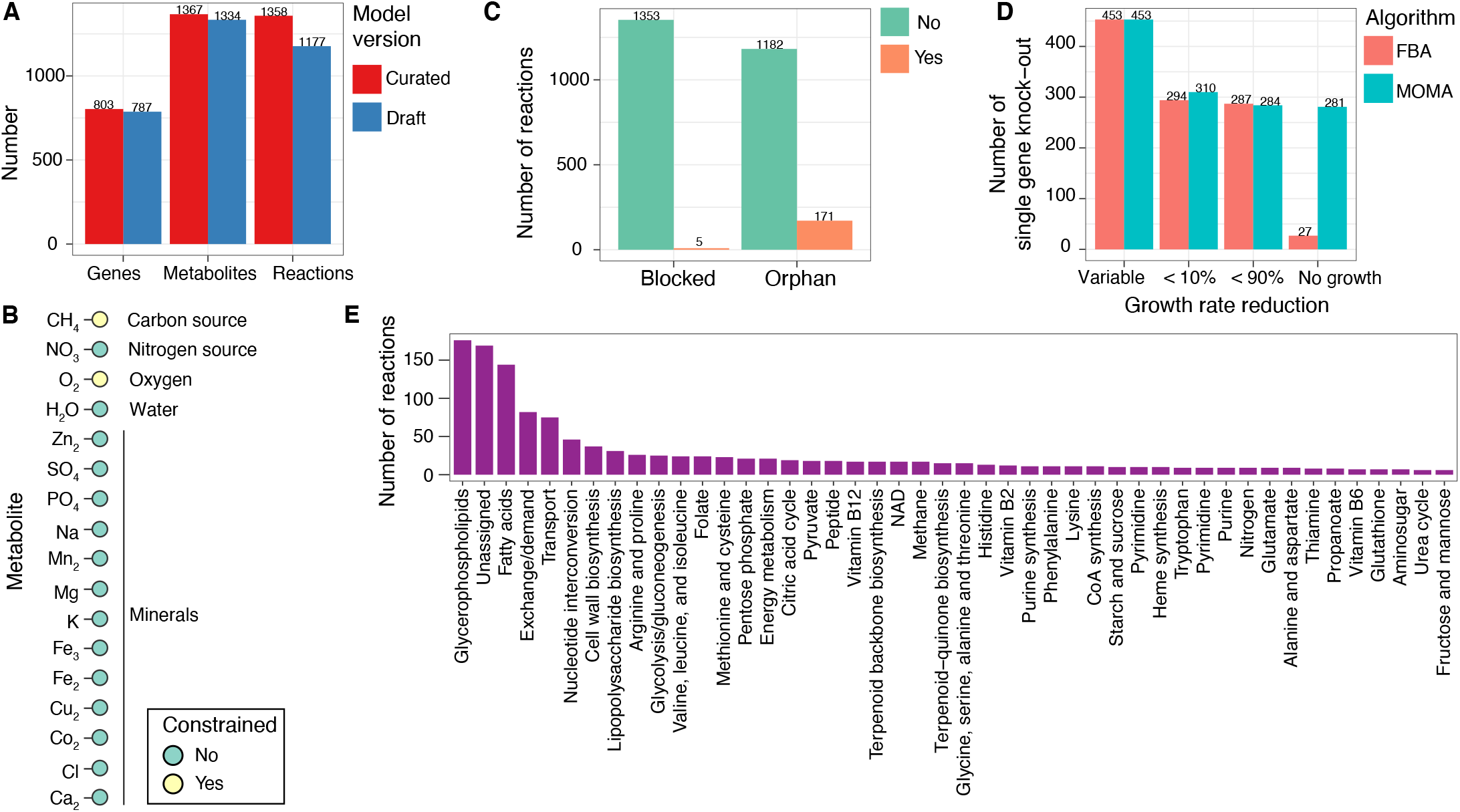
Features of the genome-scale metabolic model (GEM) of *Methylomicrobium album* BG8. **(A)** Comparison of the key features between the draft and curated models. **(B)** The default *in silico* medium composition used in the GEM. Only the methane and oxygen uptake fluxes were constrained by default. **(C)** Numbers of blocked and orphan reactions included in the curated model. **(D)** Predicted effects of single-gene knockouts on the growth rate. **(E)** Numbers of reactions comprising each pathway. Pathways with fewer than five reactions have been omitted to enhance visualization.

By applying the FBA ^42^ and minimization of metabolic adjustment (MOMA) ^43^ approaches to the model simulations, 281 essential reactions were identified in which a single gene knockout resulted in growth inhibition (Fig. 4D). Consequently, the highquality final version of the iJV803 GEM included 45 metabolic pathways containing more than five reactions (Fig. 4E). In descending order, the glycerophospholipid metabolism, (ii) fatty acids metabolism, (iii) exchange/demand and transport reactions, (iv) nucleotide interconversion, and (v) cell wall biosynthesis pathways contained the largest numbers of reactions (Fig. 4E). However, reactions that cannot be assigned to specific metabolic pathways accounted for a large proportion of the total reactions, and these remain to be classified in future studies (Fig. 4E).

### Strong predictive capabilities of the GEM of *M. album* BG8

We next tested the predictive modeling capabilities of the GEM for *M. album* BG8 using FBA. We first evaluated the model-predicted growth rate based solely on the experimentally derived methane specific uptake rates. The lower and upper 99% CIs of the methane-specific uptake rates derived from cultures with different initial oxygen-to-methane headspace ratios (Fig. 2E) were used as the uptake flux constraint. As shown in Fig. 5A, the predicted growth rates were similar to the experimental results, demonstrating the predictive power of the GEM. We also evaluated the model-predicted ratio between the oxygen and methane specific uptake rates that would maximize the growth rate, and found that the predicted flux ratio of ~1.5:1 matched the specific uptake rate ratios observed in the cultures with a higher initial oxygen headspace percentage (Fig. 5B).

**Figure 5.**
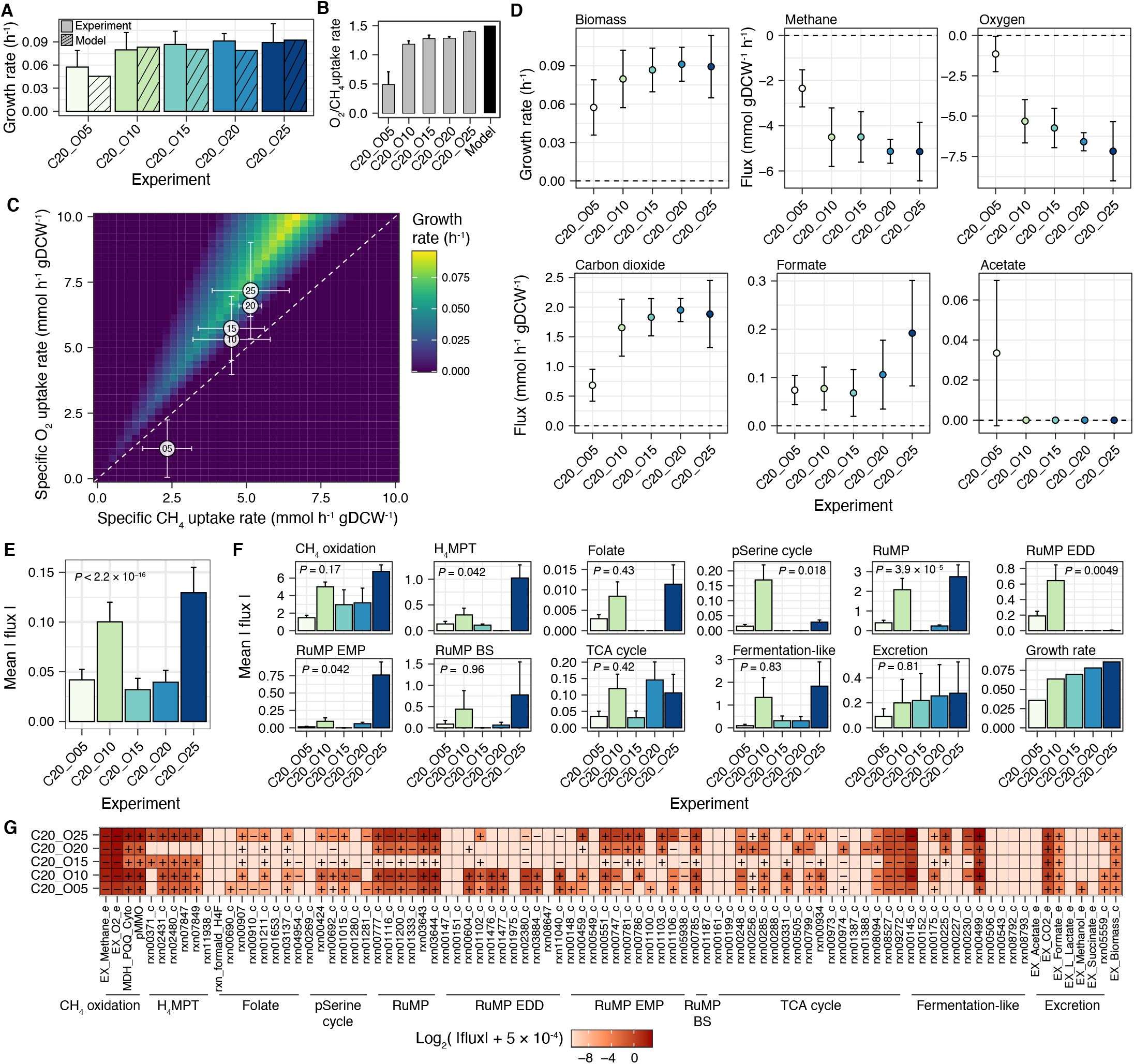
Predictive capabilities of the genome-scale metabolic model (GEM) of *Methylomicrobium album* BG8 and construc-tion of context-specific GEMs (csGEMs) under different initial percentages of oxygen (5% to 25%) in 5% increments with 20% methane. **(A)** Comparison of the experimental and predicted growth rates. Error bars on the experimental data represent the 99% confidence intervals. **(B)** Comparison of the experimental specific uptake rate ratios with the predicted optimal oxygen-to-methane uptake flux ratios. **(C)** Metabolic robustness analysis of the growth rates at different oxygen-to-methane uptake flux ratios. The white circles denote the experimentally determined specific uptake rates, and the error bars represent the 99% confidence intervals. The number inside each circle indicates the initial oxygen headspace percentage in each experiment. **(D)** Experimentally determined growth rate and fluxes; the error bars represent the 99% confidence intervals indicating statistical significance. **(E)** Mean absolute flux throughout the metabolic network in the csGEM of each experiment. **(F)** Mean absolute flux of each central metabolic pathway indicated in Fig. 3. For panels **E** and **F**, the mean values of the fluxes are shown, the error bars represent the standard errors, and the *P*-values were calculated using the Kruskal-Wallis test. **(G)** Heatmap of the flux through each reaction in the central metabolic pathways. The “-” and “+” symbols denote the directions of the reactions in the biochemical network. For each reaction, 5 × 10^−4^ *mmol h*^−1^ *gDCW*^−1^ was added to the absolute flux value to avoid zero flux and enable visualization of the flux values on a log_2_ scale.

In the same experiment (Fig. 2), the initial oxygen-to-methane headspace ratio and the specific oxygen and methane uptake rates strongly influenced the growth rates and the final excreted metabolite yields and concentrations. To investigate the variations in growth rate under different oxygen-to-methane uptake flux ratios, a robustness analysis was performed while controlling the oxygen and methane uptake flux reactions and maximizing the biomass production reaction (Fig. 5C). Consistent with the experimental results (Fig. 2), the GEM predicted a range of ratios of oxygen-to-methane uptake flux within which growth was feasible (above the slope = 1 line on Fig. 5C). The highest growth rate was predicted at a slope of ~1.5 (Fig. 5C), indicating that although biomass production may be possible at an oxygen-to-methane uptake flux ratio > 1:1, the ideal oxygen uptake flux is ~1.5 times that of methane (Fig. 5C). Interestingly, most of the experimentally determined specific uptake rate ratios (Fig. 2F) were within the predicted range of feasible growth; the experiment with the lowest initial oxygen headspace percentage (5%) was the only exception (Fig. 5C). This result suggests that the specific oxygen and methane uptake rates may not be optimal for *M. album* BG8 growth when oxygen availability is very low.

### GEM revealed the role of oxygen availability in balancing formaldehyde oxidization and assimilation

Given the apparent effect of oxygen on methane oxidation and subsequent biomass production and metabolite excretion (Fig. 2), the GEM was used to further investigate the effects of oxygen on system-level metabolism in *M. album* BG8. Using time-series data from the experiments conducted under varying initial oxygen headspace percentages (Fig. 2), the growth rates, methane and oxygen uptake fluxes, and carbon dioxide, formate, and acetate excretion fluxes in the exponential phase were computed (Fig. 5D). The growth rate increased proportionally to the initial oxygen headspace percentage (and the initial oxygen-to-methane headspace ratio) up to a maximum of 15% oxygen (Fig. 5D). The methane and oxygen uptake fluxes and carbon dioxide excretion flux were significantly different in the culture with the lowest oxygen-to-methane headspace ratio compared to the rest (Fig. 5D). Similar to the experimental results (Fig. S2), no direct correlation was observed between the formate excretion fluxes and the initial oxygen headspace percentage in the model (Fig. 5D). Nevertheless, the experiment with the highest initial oxygen-to-methane headspace ratio exhibited the highest formate excretion flux value (Fig. 5D). Acetate flux was only detected in the culture with the lowest initial oxygen-to-methane headspace ratio (albeit the concentration was low and the uncertainty was high; Fig. 5D).

The use of the upper and lower bounds of the measured fluxes as constraints on the respective reactions in the GEM enabled the construction of a csGEM for each experimental condition. Subsequently, the distribution of flux in the key central metabolic pathways was analyzed in detail (Fig. 5E to 5G). The csGEMs revealed significant differences in mean flux between the five oxygen conditions (Kruskal-Wallis test, *P* < 2.2 × 10^−16^, Fig. 5E), with the highest values predicted in the cultures with initial oxygen headspace percentages of 25% and 10% (Fig. 5E). Significant differences (Kruskal-Wallis test, *P* < 0.05) were observed in the H_4_MPT pathway, the partial serine cycle, the RuMP cycle, the RuMP EDD-variant, and the RuMP EMP-variant (Fig. 5F). Notable formaldehyde dehydrogenase (FDH, rxn00371_c, Fig. 3) activity was detected in the experiments initially provided with 15% and 25% oxygen (Fig. 5G), while the NADH-consuming glycerate production reaction (rxn01280_c, Fig. 3) was only highly active in the culture initially provided with 10% oxygen (Fig. 5G).

Overall, the results of the csGEM-based system-level analysis of *M. album* BG8 cultures provided with different initial oxygen headspace percentages reveal that these phenotypic variants differ mainly in the regulation of formaldehyde oxidization and assimilation. At an initial oxygen-to-methane headspace ratio of > 1:1, formaldehyde can be efficiently oxidized through the H_4_MPT pathway and assimilated mainly through the RuMP cycle and its EMP variant (Fig. 3 and Fig. 5F-5G). However, at an initial oxygen-to-methane headspace ratio of < 0.75:1, the folate metabolism pathway contributes to the oxidation of formaldehyde, which is then assimilated through the EDD variant of the RuMP cycle (Fig. 3 and Fig. 5F-5G). Interestingly, the model predicted that some methanol must be excreted to perform efficient methane oxidation under the condition of low initial oxygen availability (5%) (Fig. 5G). Furthermore, the culture with the highest biomass yield (20% methane and 20% oxygen) was predicted to carry a relatively lower mean absolute flux through its metabolic network (Fig. 5E) and high activity throughout the tricarboxylic acid (TCA) cycle (Fig. 3 and Fig. 5G).

### Regulation of organic compound excretion by the oxygen-to-methane uptake flux ratio

Because the initial oxygen headspace percentage had a system-level effect on the metabolic state of *M. album* BG8, a potential mechanistic link to organic compound excretion and biomass production was further investigated. Because the GEM cannot account for absolute concentrations, we focused our analysis on the oxygen to methane uptake flux ratio. The GEM was used to perform over 700 simulations in which the lower and upper bounds of the methane uptake flux were set to an arbitrary value of 10 *mmol h*^−1^*gDCW*^−1^ and the lower and upper bounds of the oxygen uptake flux were set to be the same and allowed to vary from 0 to 30 *mmol h*^−1^gDCW ^−1^. The large volume of simulation results provided insight into the effects of different combinations of methane and oxygen uptake fluxes on the production of biomass and ATP and the excretion of organic compounds, including carbon dioxide, formate, acetate, lactate, and succinate.

Oxygen-to-methane uptake flux ratios in the range of > 0:1 to < 2.5:1 were predicted to feasibly support biomass and ATP production (Fig. 6A) and organic compound excretion (Fig. 6B). The optimal ratio for biomass and ATP production was ~1.5:1 (Fig. 6A). At high oxygen-to-methane uptake flux ratios (> 2:1), carbon dioxide and formate were preferentially excreted (Fig. 6B), consistent with the experimental results measured in the exponential growth phase (Fig.1A and 2A). By limiting the analysis to excreted metabolites that could be experimentally detected (i.e., carbon dioxide, formate, and acetate), the simulation demonstrated that when the oxygen uptake flux decreased (i.e., a lower oxygen-to-methane uptake flux ratio), formate and (eventually) acetate accounted for a larger fraction of the total excreted flux, whereas the carbon dioxide excretion flux was reduced (Fig. 6C). These results are consistent with the experimental phenotypes measured during the late-exponential and stationary phases (Fig. 1A and 2A).

**Figure 6.**
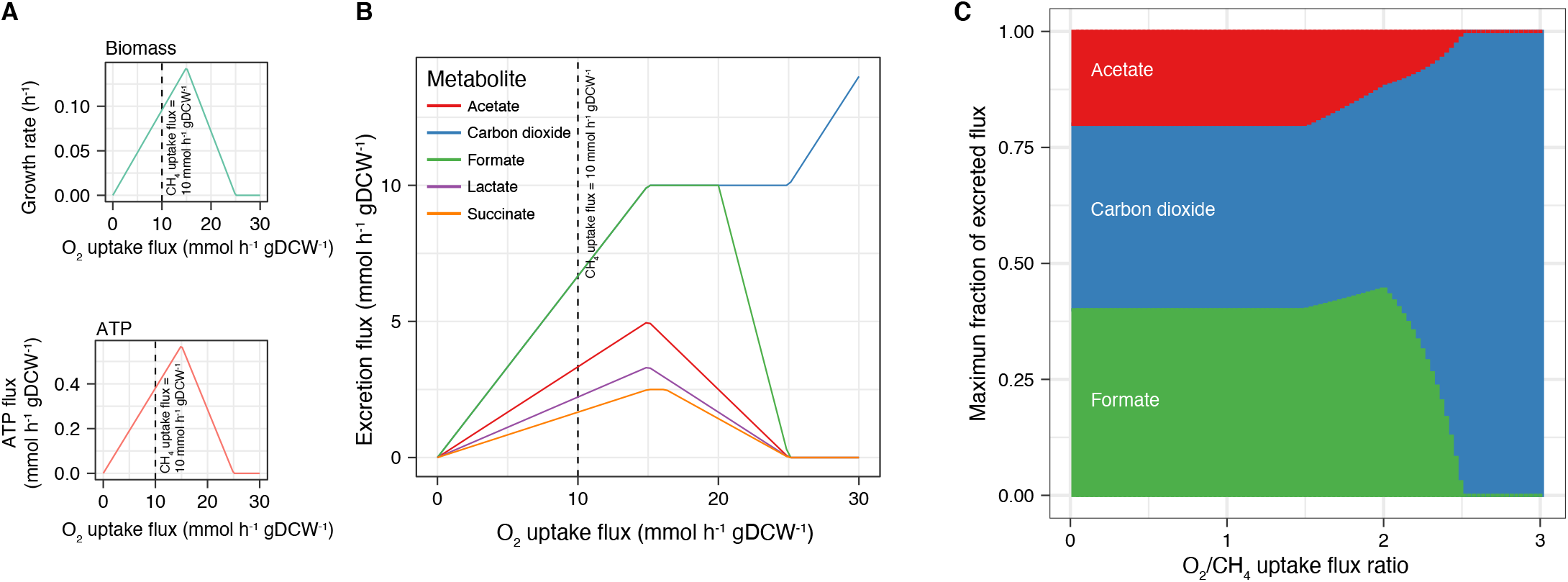
Modeled effects of the oxygen-to-methane uptake flux ratio on metabolism in *Methylomicrobium album* BG8. Effects of variations in the oxygen uptake flux on **(A)** the growth rate, ATP generation, and **(B)** organic compound excretion. Each metabolite was analyzed separately, and the excretion reaction of each was set as the objective function in the FBA. For panel **B**, the carbon dioxide and formate lines partially overlap. **(C)** Analysis of the maximum fractions of excreted flux for the three metabolites detected experimentally at different oxygen-to-methane uptake flux ratios. A constant methane uptake flux of 10 *mmol h*^−1^ *gDCW*^−1^ was used in all of the analyses.

### Metabolic modeling of a high acetate-excreting phenotype identified a mixed mode involving both respiration and fermentation

Studies ^44–48^ have determined that the initial biomass concentration can affect microbial growth and metabolic capabilities. Here, we further investigated the effects of two relatively different initial biomass concentrations (low and high) on system-level metabolism in *M. album* BG8. Both biomass concentrations were tested under the initial oxygen-to-methane headspace ratios that promoted the highest biomass and organic compound excretion yields [1:1 (20% methane and 20% oxygen) and 1.25:1 (20% methane and 25% oxygen)]. Under both ratios, the initial biomass concentration had a significant (*P* < 0.05) and specific effect on the formate and acetate yields (Fig. 7A). In the cultures with a lower initial biomass concentration, a higher formate yield was achieved under the 1:1 oxygen-to-methane headspace ratio compared with the 1.25:1 ratio (Fig. 7A). However, the acetate yield was higher in the cultures with a relatively higher initial biomass concentration under both oxygen-to-methane headspace ratios (Fig. 7A).

**Figure 7.**
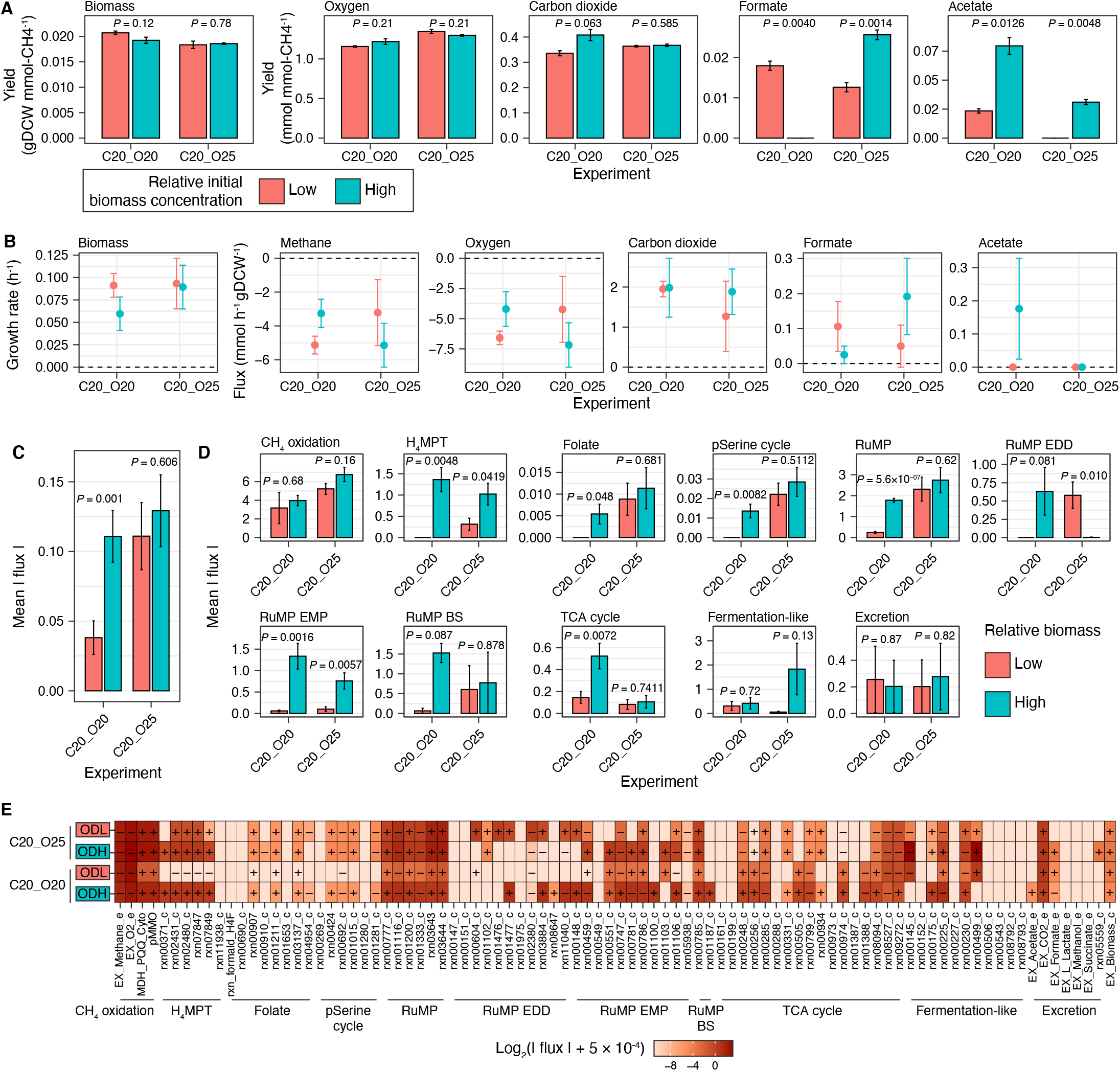
Effects of the initial biomass concentration on metabolism in *Methylomicrobium album* BG8 under two initial oxygen-to-methane headspace ratios (20% methane and 20% oxygen; 20% methane and 25% oxygen). **(A)** Biomass, oxygen uptake, and excreted metabolite yields. The data are shown as mean values of biological triplicates, and the error bars represent the standard errors. **(B)** Experimentally determined flux values; the error bars represent the 99% confidence intervals. **(C)** The mean absolute flux throughout the metabolic network in the context-specific genome-scale metabolic model (csGEM) of each experiment. **(D)** The mean absolute flux in each central metabolic pathway indicated in Fig. 3. For panels **C** and **D**, the mean values of the fluxes are shown, the error bars represent the standard errors, and the *P*-values were calculated using the two-sided *t*-test. **(E)** Heatmap of the flux through each reaction in the central metabolic pathways. ODL and ODH represent low and high relative initial biomass concentrations, respectively. The “-” and “+” symbols denote the directions of the reactions in the biochemical network. In each reaction, a value of 5 × 10^−4^ *mmol h*^−1^ *gDCW*^−1^ was added to the absolute flux value to avoid a zero flux and facilitate visualization of the flux values on a log_2_ scale.

No consistent trends in the methane and oxygen uptake fluxes were observed between the cultures with high and low initial biomass concentrations grown under different initial oxygen-to-methane headspace ratios (zFig. 7B). Furthermore, the biomass concentration had a significant effect on formate and acetate excretion (Fig. 7B, statistical significance was determined by the 99% confidence intervals represented by the error bars) but not on carbon dioxide excretion. Acetate excretion flux was observed during the exponential phase under a relatively higher initial biomass concentration and an initial oxygen-to-methane headspace ratio of 1:1 (Fig. 7B). However, these conditions led to a significantly lower growth rate compared to the other three cultures (Fig. 7B).

Using csGEMs to model the metabolism of *M. album* BG8 revealed that only the culture with a relatively higher initial biomass concentration and an initial oxygen-to-methane headspace ratio of 1:1 exhibited significantly higher (*t*-test, *P* < 0.05) mean absolute fluxes in the overall metabolic network (Fig. 7C) and in several metabolic pathways, including H_4_MPT, folate metabolism, the partial serine cycle, RuMP cycle, the EMP variant of the RuMP cycle, and the TCA cycle (Fig. 7D). In the cultures grown under an initial oxygen-to-methane headspace ratio of 1.25:1, we identified significant effects of the initial biomass concentration (*P* < 0.05) on the mean absolute fluxes in the metabolic pathways of H_4_MPT and the EMP and EDD variants of the RuMP cycle (Fig. 7D).

A detailed analysis of affected pathways and reactions in the acetate-excreting phenotype indicated that the BS variant of the RuMP cycle, the reactions of which are linked to fermentation-like reactions, exhibited a high level of flux (Fig. 7E). In fact, high levels of flux (Fig. 7E) were observed in the reaction that converts fructose-6P plus glyceraldehyde-3P to xylulose-5P and erythrose-4P (rxn00785_c, Fig. 3), and the subsequent reaction that converts xylulose-5P to acetyl-P (rxn01187_c, Fig. 3). Acetyl-P and acetate can be interconverted through the reactions rxn00230_c and rxn00225_c (with ATP generation), both of which exhibited high levels of flux in this phenotype (Fig. 7E). Furthermore, the csGEM of this acetate-excreting phenotype predicted a second pathway leading to acetate excretion: specifically, high flux was observed in the pathway that converts pyruvate to acetyl-CoA (rxn05938_c) and acetyl-CoA to acetate (rxn00175_c) (Fig. 7E and Fig. 3) and also generates ATP (Fig. 3). Intriguingly, the reactions that convert malate to malyl-CoA (rxn00934, Fig. 3) and malyl-CoA to acetyl-CoA plus glyoxalate (rxn00331_c, Fig. 3) both exhibited high levels of flux, suggesting that this pathway might replenish acetyl-CoA at the expense of ATP and connect the TCA cycle with the partial serine cycle. In turn, the active production of malate and subsequent generation of acetyl-CoA would explain the high flux in the TCA cycle in this phenotype (Fig. 7E). Interestingly, the flux analysis of this acetate-excreting phenotype also revealed that the entire TCA cycle was used to produce one molecule of ATP and two molecules of NADH and to excrete carbon dioxide (Fig. 7E and Fig. 3). Taken together, the results of integrative modeling in this study suggest that the high acetate-excreting phenotype uses a mixed metabolic mode in which the pathways required for respiration and fermentation are simultaneously active.

### Metabolic versatility of *M. album* BG8 and trade-offs between biomass production and organic compound excretion

After separately analyzing and describing the effects of the initial methane (Fig. 1), oxygen (Fig. 2), and biomass (Fig. 7) concentrations on the metabolism of *M. album* BG8, we quantitatively investigated the associations of these factors with the oxygen-to-methane uptake flux ratio, an important parameter used to define phenotypes (Fig. 6). In all of the experiments, the relationship between the initial oxygen-to-methane headspace ratio and the calculated oxygen-to-methane uptake flux ratio could be described using a bounded exponential model (Fig. 8A). At an initial oxygen-to-methane headspace ratio > 0.5:1, the maximum oxygen-to-methane uptake flux ratio was 1.3:1 to 1.4:1 (Fig. 8A), consistent with the GEM-predicted optimal oxygen-to-methane uptake flux ratio of ~1.5:1 for growth and ATP generation (Fig. 6A).

**Figure 8.**
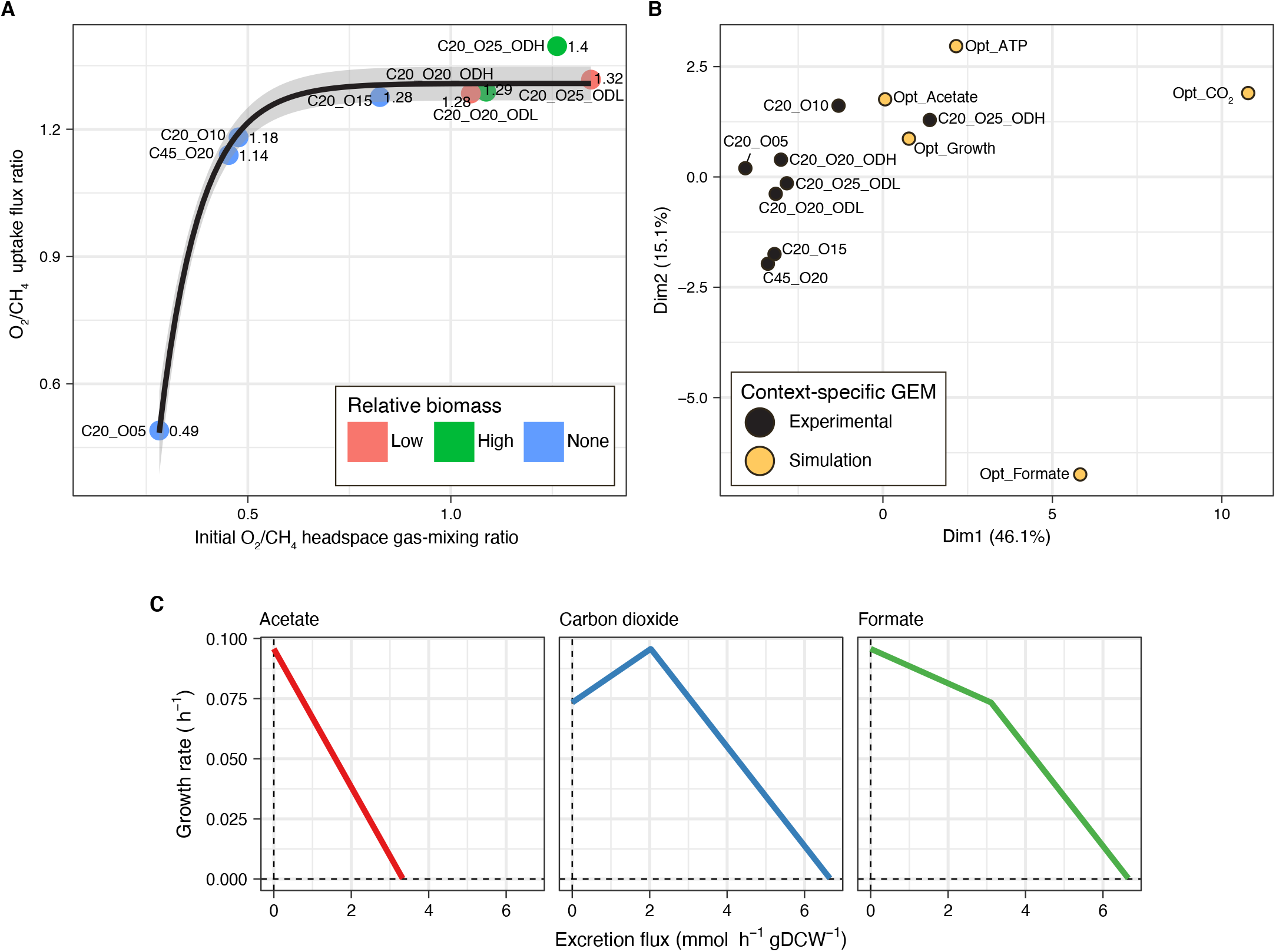
Metabolic versatility of and metabolic trade-offs in *Methylomicrobium album* BG8. **(A)** The oxygen-to-methane uptake flux ratio as a function of the initial oxygen-to-methane headspace ratio under all of the experimental conditions. A bounded exponential model was fitted to the data; the gray-shaded region represents the 95% confidence interval. **(B)** Results of a principal component analysis (PCA) of the overall metabolic flux in each experimental culture. Five simulated context-specific genome-scale metabolic models (csGEMs) were included in the PCA to represent phenotypes optimized for growth, ATP generation, carbon dioxide production, formate excretion, or acetate excretion. **(C)** Metabolic robustness analysis wherein the acetate, carbon dioxide, or formate excretion reaction was controlled and the biomass production reaction was maximized. The lines indicate the trade-off between the rate of growth and the excretion flux of a metabolite. Five hundred simulations were performed for each metabolite.

A PCA was applied to the results of the csGEM to compare the metabolic flux between different cultures (Fig. 8B). Based on the results, most of the experimental phenotypes were clustered together, except the phenotype grown under 20% methane, 20% oxygen, and a relatively high biomass (C20_O20_ODH) (Fig. 8B). This phenotype also yielded an oxygen-to-methane uptake flux ratio of 1.4:1, which deviated from the confidence interval of the bounded exponential model (Fig. 8A). Cultures with different methane and oxygen headspace percentages but similar initial oxygen-to-methane headspace ratios also had similar oxygen-to-methane uptake flux ratios (Fig. 8A) but different metabolic fluxes, as indicated by their separation on the PCA plot (Fig. 8B).

The results of csGEM simulations of phenotypes optimized for growth, ATP generation, or carbon dioxide, formate, and acetate excretion were also included in the PCA as reference values (Fig. 8B). Although the predicted metabolic fluxes tended to differ between the experimental cultures, these values were relatively closer to the reference phenotypes optimized for growth, acetate excretion, and ATP generation and clearly distant from the phenotypes optimized for carbon dioxide and formate excretion (Fig. 8B). Possibly, this metabolic versatility enabled *M. album* BG8 to optimize for growth and ATP generation under different conditions by using different pathways.

An analysis of the metabolic fluxes in the optimal phenotype for acetate production revealed a cluster that included the phenotypes optimized for growth and ATP generation and the predicted phenotypes of the experimental cultures. Possibly, acetate is a highly relevant metabolite that balances the pathways contributing to energy and biomass production in *M. album* BG8. In fact, we hypothesized that a trade-off existed between biomass production and acetate excretion in this strain. This hypothesis was partially supported by the observation of reduction in the growth rate when acetate was excreted during the exponential phase (Fig. 7B).

To test our hypothesis, we applied a robustness analysis in which the acetate excretion reaction was controlled and the biomass production reaction was maximized. The lower and upper bounds of both the methane and oxygen uptake fluxes were set to −10 *mmol h*^−1^ *gDCW*^−1^ and 0 *mmol h*^−1^ *gDCW*^−1^, respectively. This enabled the GEM to automatically adjust the oxygen-to-methane uptake flux ratio to maximize the growth rate under every acetate excretion condition. Consistent with the hypothesis, the robustness analysis revealed a trade-off between the growth rate and acetate production (Fig. 8C). However, this analysis also revealed different trade-offs between the growth rate and carbon dioxide and formate excretion (Fig. 8C). In fact, a carbon dioxide excretion flux between 0 and 2 *mmol h*^−1^ *gDCW*^−1^ (Fig. 8C), which matched the experimentally measured range, favored the growth rate (Fig. 5D and 7B).

## Discussion

The aim of this study was to elucidate the system-level metabolism Although carbon dioxide and formate are usually excreted of *M. album* BG8 using an integrative systems biology framework. The integration of time-series exometabolomics data into a newly developed, high-quality GEM yielded novel insights into the metabolic mechanisms of this methanotrophic bacterial strain. Furthermore, the experimental data collected in this study enabled the construction of csGEMs, and the application of an FBA enabled the first predictions of the system-level metabolism of *M. album* BG8 under different initial concentrations of oxygen, methane, and biomass.

The conversion factor of 0.26 *gDCW l*^−1^ derived between the *OD* 600 and DCW of *M. album* BG8 in our study differs from a previously reported conversion factor of 0.33 *gDCW l*^−1^ ^49^. Although a linear regression of our data with a y-intercept of −0.044 yielded a conversion factor of 0.32 *gDCW l*^−1^ (Fig. S1), this model was not used due to a lower adjusted *R*^2^ and a less significant *P* value compared to our selected model. These results highlight the need for caution regarding potential variability in the conversion factor between *OD*_600_ and DCW when studying *M. album* BG8. However, the Monod model was used to estimate growth kinetics as a function of the initial oxygen headspace percentage, and the calculated *μ_max_* value (0.11 *h*^−1^) is similar to other reported growth rates for *M. album* BG8 (e.g., 0.10 *h*^−1^ to 0.18 *h*^−1^ with different concentrations of methane ^24,49^, 0.09 *h*^−1^ to 0.13 *h*^−1^ with different concentrations of chloromethane ^50^, and 0.11 *h*^−1^ with methanol ^51^). However, the Monod model assumed that oxygen was the only limiting substrate, which may not be true in a real-life scenario.

We identified oxygen as the key driver of methane oxidation by *M. album* BG8, as it exerted strong effects on both biomass production and organic compound excretion. In our analysis, optimal growth could be sustained at a ratio of > 1:1 between the specific uptake rates of oxygen and methane. Biomass production was maximized under culture conditions of 20% methane and 20% oxygen, with a yield of 0.021 *gDCW mmolCH*4^−1^. This was the only phenotype variant predicted by the csGEM to have a completely active TCA cycle and to exhibit a low mean level of absolute flux throughout its metabolic network. Together, these results of integrative *M. album* BG8 modeling indicate an optimal methanotrophic state for the allocation of molecular and metabolic resources in which optimal biomass production is preserved and enzyme use is minimized ^52^. These results also complement the findings of recent work in which *M. album* BG8 was identified as the bacterium with the highest biomass yield (nearly double that of the others tested) among a group of industrially relevant methanotrophs ^25^.

Intriguingly, the GEM predicted the excretion of methanol as a by-product of growth under the condition with the lowest initial oxygen availability (5%). A possible explanation for this predicted phenotype is that methanol works as an additional control for the metabolic branching at low oxygen concentration, leading to tetrahydrofuran-based oxidation of formaldehyde in addition to metabolic flux through the EDD variant of the RuMP cycle and partial serine pathways. Hence, instead of actual methanol release to the extracellular environment (which was not detected in the medium), *M. album* BG8 could be using the mentioned pathways to more efficiently oxidize methanol to gain biomass. A recent transcriptomics and metabolomics study of *M. album* BG8 grown on methanol ^53^ yielded a phenotype that is similar to the condition with low availability of oxygen. Increased transcription of genes related to carbon metabolism through the RuMP EDD-variant and pentose phosphate pathways as well as formaldehyde detoxification through the glutathione dependent pathway, which formate is the product, were observed ^53^. Taking the reported transcriptomics and metabolomics data ^53^ together with the GEM and FBA results presented here, they suggest that *M. album* BG8 potentially favors methanol oxidation over methane as oxygen availability diminishes.

Although carbon dioxide and formate are usually excreted during aerobic methanotrophy ^54^, these metabolites have recently become valuable by-products ^55,56^ and are used as carbon sources in synthetically constructed methanotrophic modular microbial consortia to produce value-added compounds ^57^. In our study, the highest carbon dioxide yield (0.39 *mmol mmolCH*4^−1^) was achieved under culture conditions of 20% methane and 20% oxygen, and the highest formate yield (0.025 *mmol mmolCH*4^−1^) was obtained under conditions of 20% methane and 25% oxygen. In contrast to a recent report ^25^, formate excretion was detected in all of the cultures in this study. The phenotype with the highest formate yield exhibited the highest mean absolute flux through its metabolic network and the second highest flux through the H_4_MPT formaldehyde oxidation pathway. In other words, by actively using the complete H_4_MPT pathway, this phenotype can generate two reducing equivalents (NAD(P)H). Based on these results, developers of metabolic engineering strategies that aim to optimize carbon dioxide and/or formate excretion by *M. album* BG8 during methane metabolism may consider directing carbon flux toward the H4MPT pathway. Nevertheless, future attempts to optimize formate production by *M. album* BG8 should consider the achievement of significantly larger formate yields by other methanotrophs grown on methanol instead of methane ^58–60^. Although a previous study investigated the effects of methane and methanol on formate excretion by *M. album* BG8, the authors did not detect formate under the tested conditions ^25^, likely because the samples collected in the reported experiments corresponded to the early stages of *M. album* BG8 growth in our experimental setup.

Acetate is a model metabolite derived from acetyl-CoA ^61^, a key precursor in high-value chemical production ^62^. Therefore, the biotechnological potential of *M. album* BG8 in the conversion of methane to valuable compounds depends on the ability to produce a high acetate yield. Among the different culture conditions in this study, the highest acetate yield (0.072 *mmol mmolCH*4^−1^) was unexpectedly obtained under the conditions of 20% methane, 20% oxygen, and a relatively high initial biomass concentration. This phenotypic variant of *M. album* BG8 excreted acetate during the exponential phase, demonstrating the potential coupling of exponential growth with acetate production in this strain. Interestingly, a trade-off between this high acetate yield and biomass production occurred, resulting in a lower growth rate. However, the molecular and metabolic mechanisms behind this trade-off remain unclear and need to be further investigated. Previous reports have described acetate excretion by other gammaproteobacterial methanotrophs under prolonged oxygen starvation ^63^ and oxygen-limited growth ^64^. Our experimental results agree with those findings, as acetate excretion was detected in all cultures in which oxygen was present at a very low concentration or completely depleted. Similarly, our model predicted a high acetate excretion flux in *M. album* BG8 at a low oxygen-to-methane uptake flux ratio. Overall, we expect that the metabolic characterization of acetate excretion presented herein will support future attempts to increase acetate yields and achieve the metabolic reprogramming of acetyl-CoA conversion in *M. album* BG8.

Both the oxygen-to-methane headspace ratio and specific uptake ratio have been explored intensively in methanotrophy studies ^58,63,65–68^, as these variables are thought to control the differential activation of pathways required for the production of biomass, generation of energy, and induction of fermentation-like metabolism ^63,64^. The *M. album* BG8 genome encodes homologs of enzymes found in the EDD and EMP RuMP cycle variants in *M. alcaliphilum* 20Z and *M. buryatense* 5G(B1) ^64^. However, *M. album* BG8 lacks 3 of the 18 key enzymes (i.e., phosphate acetyltransferase, D-fructose 6-phosphate phosphoketolase, and NAD(P)-dependent malic enzyme) required for fermentation-like metabolism in *M. alcaliphilum* 20Z ^64^ and *M. buryatense* 5G(B1) ^63^ at low oxygen-to-methane concentration ratios. Nevertheless, the *M. album* BG8 cultures exhibited fermentation-like phenotypes with high acetate excretion and low carbon dioxide and formate excretion, mainly during the stationary phase when the oxygen-to-methane concentration ratio was very low. Finally, we identified a non-linear relationship between the initial oxygen-to-methane headspace ratio and the oxygen-to-methane uptake flux ratio in batch cultures of *M. album* BG8, which will facilitate control of the oxygen-to-methane uptake flux ratio in batch systems

In summary, the experimental results and the metabolic flux predictions presented in this study elucidate some of the characteristics of *M. album* BG8. However, many other aspects of metabolism in this strain remain to be explored in detail. The newly developed GEM of *M. album* BG8 is a valuable tool that will aid future investigations of this important methanotroph in the context of methane conversion applications in biomanufacturing or methane emission mitigation in environments that experience dynamic fluxes of methane and oxygen.

## Methods

### Strain

*M. album* BG8 was purchased from American Type Culture Collection (USA; catalog number 33003DTM). The genome of this strain has been sequenced ^69^ and is publicly available in the National Center for Biotechnology Information (NCBI) repository (assembly: ASM21427v3), the Integrated Microbial Genomes (IMG) system (IMG Genome ID: 2508501010), and the KBase Central Store (Source ID: 686340.3).

### Experimental procedures

For all of the experiments, *M. album* BG8 was cultured in 20 mL of nitrate mineral salts (NMS) medium ^70^ in a 160-mL serum bottle capped with a butyl rubber stopper. The initial bacterial cell concentration was adjusted according to the experimental specifications. The culture bottles were incubated at 30 °C with agitation at 200 rpm, and CuCl2 (5 μM) was used as the copper source ^71^. The headspace was filled with high-purity (99.995%) nitrogen, and the desired initial mixing ratio between oxygen and methane in the headspace was obtained by injecting appropriate volumes of the respective high-purity (99.7%) gases with a syringe. Here, the initial headspace volume-per-volume percentage refers to the reported percentages of oxygen and methane. All of the experiments were performed with three biological replicates. Uninoculated controls were set up in parallel to assess abiotic losses (which were insignificant). For downstream analyses, the cells were collected on 0.22-μm filters (47 mm, MF-Millipore^TM^, MilliporeSigma, USA) and washed twice with sterile NMS medium.

### Analytical analysis

To determine the headspace concentrations (mM) of hydrogen, oxygen, methane, and carbon dioxide, samples were collected using a gas-tight syringe (Hamilton, USA) and analyzed on a gas chromatograph (GC-2010, Shimadzu, Japan) as described previously ^72^. Excreted metabolites in the aqueous phase were measured using a high-performance liquid chromatography (HPLC) system (Waters Corporation, USA) equipped with an Aminex Ion Exclusion HPX-87H column (Bio-Rad, USA) as described previously ^72^.

The concentrations of cellular amino acids and total lipid fatty acids in bacterial cells were analyzed during the exponential growth phase. Seventeen amino acids were measured quantitatively using HPLC at the Proteomics and Metabolomics Facility, Center for Biotechnology, University of Nebraska-Lincoln (USA) following standard protocols. The total levels of 15 lipid fatty acids were measured quantitatively using thin-layer chromatography, which was performed by a commercial laboratory (AminoAcids.com, USA) following standard protocols.

### Conversion between the optical density and dry cell weight

To determine the conversion factor between the observed optical density at 600 nm (OD_600_) and the dry cell weight (DCW), 13 cultures of *M. album* BG8 were grown under 20% methane and 20% oxygen to varying cell densities (OD_600_ 0.2 to 1.0), and the corresponding lyophilized cell masses were weighed. OD was meaured using a UV-VIS spectrophotometer (UV-1800, Shimadzu, Japan), and DCW was determined using an MS-TS analytical balance (MS204, Mettler Toledo, USA).

### Calculations of yield, rate, and flux

The yields of total biomass (*gDCW mmolCH*4^−1^) and of specific metabolites (*mmol mmolCH*4^−1^) were calculated by considering the changes in concentration per total methane consumed throughout the experiment. The growth rate (*h*^−1^) was calculated over the exponential phase as the log-linear regression between the biomass concentration and time. The specific uptake rate and excretion rate were considered equivalent to the uptake flux and excretion flux, respectively, when integrated as constraints into the GEM. To meet the steady-state assumption of the FBA, both rates or fluxes were calculated for each metabolite (in *mmol h*^−1^ *gDCW*^−1^) in each experiment, using a linear regression of only those data points that were collected during the exponential phase as described previously ^73^. The computed confidence intervals of the slopes in the linear regression model informed the uncertainty of the measured data and were applied as upper and lower constraints to the reactions in the GEM.

### Reconstruction of a draft GEM

The GEM for *M. album* BG8 was reconstructed based on its genome sequence ^69^. A draft version of the model was reconstructed using KBase ^41^ and converted into Systems Biology Markup Language (SBML) for further curation and processing using the Sybil package in R ^74^, the COBRApy library in Python ^75^, and the COBRA Toolbox 3.0 in MATLAB ^40^.

### *In silico* growth medium

An *in silico* recipe of the chemical compounds required to produce biomass (i.e., growth medium) must be inputted into a GEM. All of the components in the experimental NMS medium ^70^ were mapped to the KBase compounds database using the tools in the Known Media Database (KOMODO) ^76^. In the GEM, methane and nitrate were set as the sole carbon and nitrogen sources, respectively. The oxygen exchange reaction was set at a lower limit of −10 *mmol h*^−1^ *gDCW*^−1^ to ensure an aerobic growth condition.

### Integration of experimentally determined biomass compositions into the GEM

In a GEM, an accurate representation of the biomass composition enables better representations of the associated cellular biology and biochemistry. The biomass composition is specified using a metabolic reaction that accounts for all of the metabolites (and their respective concentrations) needed to generate 1 g of DCW. However, the automatically reconstructed GEM in KBase only generated a draft biomass production reaction in which the default biomass composition was generated according to the Gram classification of the bacterium of interest. To account for the experimentally determined biomass composition, the *M. album* BG8 biomass production reaction was manually edited using Python on the Jupyter Notebooks ^77^ interface of KBase. Briefly, the reaction was updated by adding the experimentally measured concentrations of amino acids and fatty acids and the intracellular organic metabolite, trace element, and cofactor concentrations used in the GEM of *M. buryatense* 5G(B1) ^32^. Pyrroloquinoline quinone (PQQ), a cofactor specific for methanol dehydrogenase, was added manually to the biomass production reaction. Other biomass components were obtained from the automatic KBase GEM reconstruction. The biomass production reaction in the GEM was then formulated using the stoichiometric coefficients of all of the metabolites required to generate 1 g of DCW. The following assumptions were made: growth-associated ATP maintenance of 59.8 *mmol gDCW*^−1^, which is used in the GEMs of *Escherichia coli* ^78^ and other methanotrophs ^32,35^, and non-growth-associated ATP maintenance of 3.5 *mmol h*^−1^ *gDCW*^−1^, as reported elsewhere for the methanotroph *M. parvus* OBBP ^37^.

### Gap-filling and model refinement

After the biomass production reaction in the GEM was edited as described in the previous section, automatic gap-filling of the draft GEM was performed using KBase and components of the *in silico* NMS medium as the nutrient source. All of the reactions added in the gap-filling step were manually checked to determine their essentiality in the GEM using a leave-one-out approach, wherein single reactions were excluded and an FBA was applied to maximize the biomass production reaction. Reactions that were not essential to *in silico* biomass production were discarded.

After gap-filling, the model was fine-tuned using an iterative refinement approach as described previously ^40,79^. The geneprotein-reaction associations, reaction directionality, co-factor and co-enzyme assignments, and Enzyme Commission numbers in the draft GEM were subjected to manual curation, using the following as references: (i) published literature, (ii) NCBI assembly annotations, (iii) BlastKOALA ^80^ genome annotation, (iv) RAST ^81^ genome annotation and the (v) KEGG ^82^, (vi) BRENDA ^83^, and KBase ^41^ reaction and metabolite databases.

The KBase-generated draft GEM in SBML format did not contain the chemical formulae of metabolites. The formulae from the KBase biochemistry database were manually integrated into the GEM. The balance of carbon, hydrogen, oxygen, and nitrogen was checked using the checkBalance function in the COBRA Toolbox 2.0. The final version of the *M. album* BG8 GEM was named **iJV803** according to standard convention 84, where **i** stands for *in silico*, **J** and **V** are the initials of the name of the GEM builder, and **803** represents the number of included metabolic genes.

### Gene locus tags in the GEM

*M. album* BG8 genes found in KBase (Source ID: 686340.3) were identified using locus tags that differed from those in the NCBI (accession: NZ_CM001475) and Joint Genome Institute (Genome ID: 2508501010) depositories. To reconcile these differences and facilitate the use of the GEM in future research applications, all of the genes present in the GEM were aligned against genes identified in different repositories using blast ^85^ (identity cutoff: 99%) to identify the corresponding locus tags. The annotations of all of the key genes involved in methane oxidation were further verified against the reported functions in ^24,69^, with no discrepancies. The locus tags of all of the genes in the GEM and their corresponding annotations in the different repositories are summarized in Table S1.

### Quality control and quality assurance

The quality of the iJV803 GEM was checked using the MEMOTE platform^86^,and a report of the model can be obtained at (https://juanvillada.github.io/iJV803/docs/index.html).

### Genome-scale metabolic modeling and FBA

FBA ^42^ was performed using the R package Sybil ^74^. To avoid infeasible metabolic cycles, the cycle-free FBA method (cfFBA) was also applied to metabolic modeling using the R package CycleFreeFlux ^87^ Linear programming problems were solved using the GNU Linear Programming Kit (version 4.65). Although the objectives of optimization varied according to the objective of each analysis ^88^, the biomass production reaction was the default objective function in the GEM. The FBA method was applied in all metabolic robustness analyses, whereas the cfFBA method was used to optimize csGEMs. After normalizing the metabolic flux of each reaction to the methane specific uptake rate of each csGEM, a principal component analysis (PCA) of the metabolic fluxes of all reactions in the csGEMs was performed using the R function prcomp.

## Data availability

The final version of the model can be downloaded in RData, SBML, JSON, and XLS formats from (https://juanvillada.github.io/iJV803).

## Supplementary Data

Supplementary Data are available online.

## Acknowledgements

JCV acknowledges support provided by the Hong Kong PhD Fellowship Scheme (HKPFS).

JCV, MFD, CKL and PKHL conceived the study. JCV built the model and developed the scripts for data analysis. MFD performed the experiments. JCV, MFD, and PKHL performed data analysis and interpreted the results. LYS contributed to the interpretation of the findings. JCV, MFD, LYS and PKHL wrote the manuscript. All authors approved the final version of the manuscript.

---

**Figure S1.**
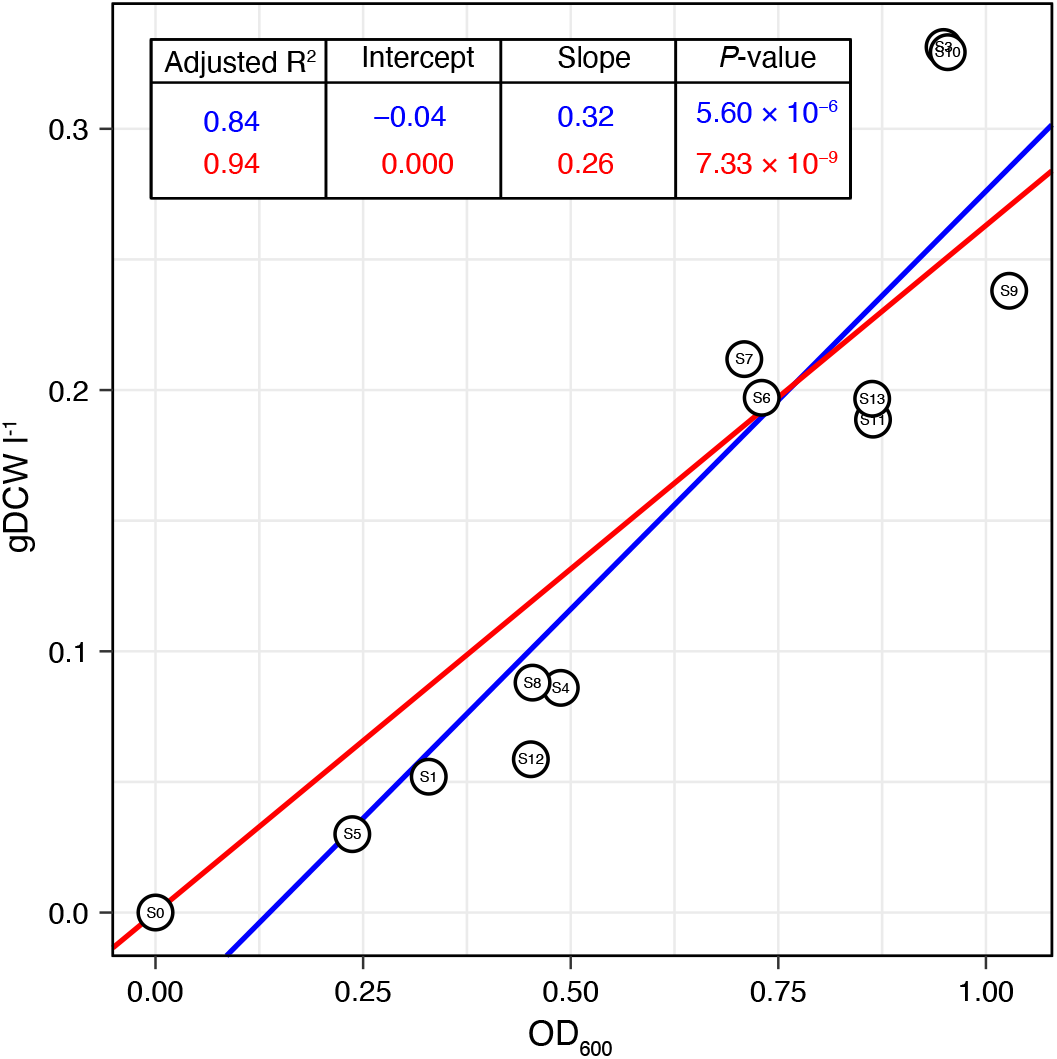
Central methanotrophic metabolic pathways in *Methylomicrobium album* BG8. Metabolites that are taken up are highlighted in yellow, and those excreted are labeled in blue. Metabolite names are shown in black, reaction identification numbers are in green, and the genes associated with each reaction are in brown. The pathway names are bolded. The chemical formula describing the key metabolites involved in the initial methane oxidation and the consumed and excreted metabolites is shown. Non-reversible and reversible reactions, determined according to the thermodynamics constraints of every reaction, are indicated by single-headed and double-headed arrows, respectively.

**Figure S2.**
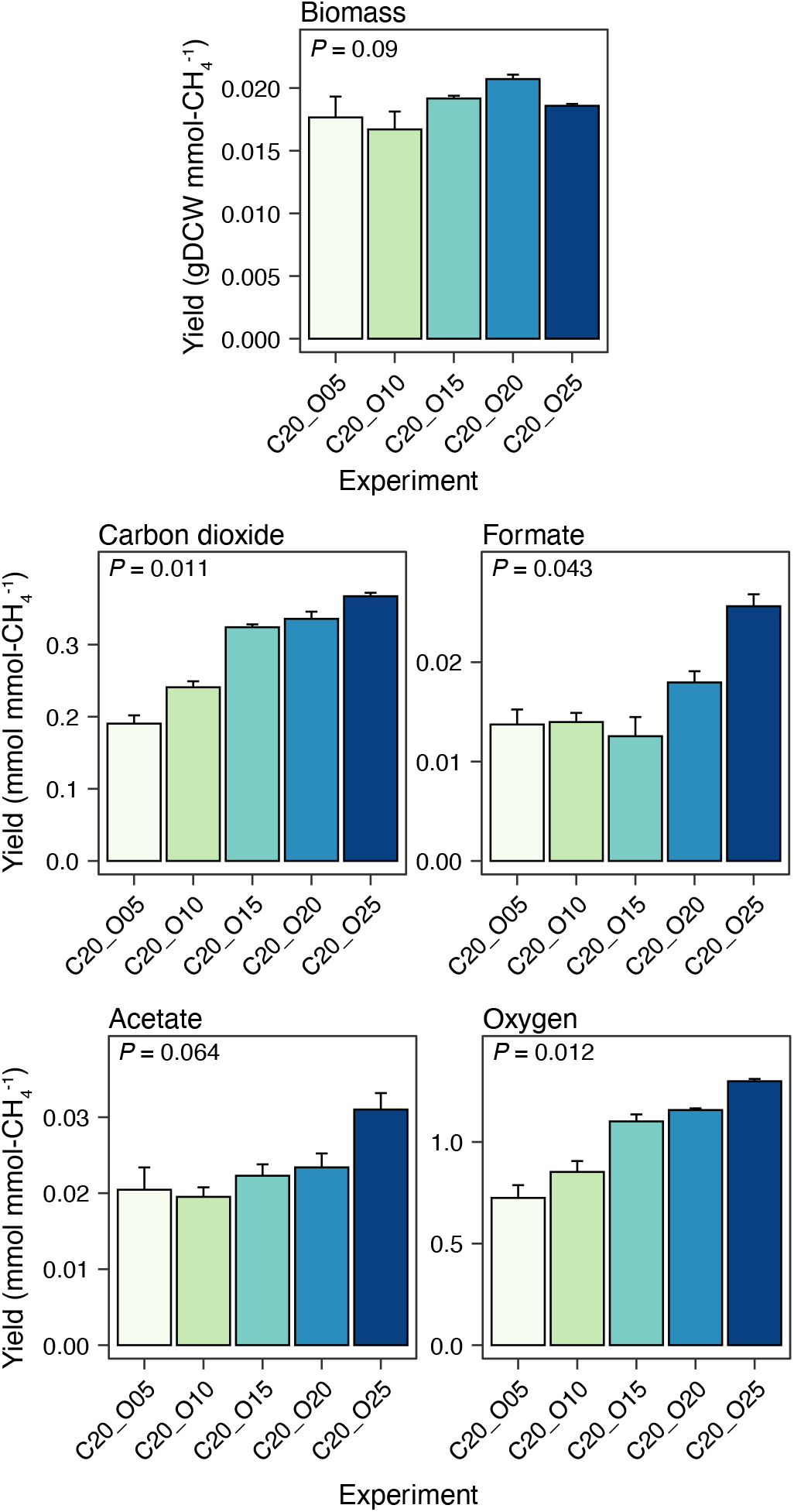
Central methanotrophic metabolic pathways in *Methylomicrobium album* BG8. Metabolites that are taken up are highlighted in yellow, and those excreted are labeled in blue. Metabolite names are shown in black, reaction identification numbers are in green, and the genes associated with each reaction are in brown. The pathway names are bolded. The chemical formula describing the key metabolites involved in the initial methane oxidation and the consumed and excreted metabolites is shown. Non-reversible and reversible reactions, determined according to the thermodynamics constraints of every reaction, are indicated by single-headed and double-headed arrows, respectively.

